# Thalamocortical dynamics underlying spontaneous transitions in beta power in Parkinsonism

**DOI:** 10.1101/422238

**Authors:** Carolina Reis, Andrew Sharott, Peter J. Magill, Bernadette van Wijk, Thomas Parr, Peter Zeidman, Karl Friston, Hayriye Cagnan

## Abstract

Parkinson’s disease (PD) is a neurodegenerative condition in which aberrant oscillatory synchronization of neuronal activity at beta frequencies (15-35 Hz) across the cortico-basal ganglia-thalamocortical circuit is associated with debilitating motor symptoms, such as bradykinesia and rigidity. Mounting evidence suggests that the magnitude of beta synchrony in the parkinsonian state fluctuates over time, but the mechanisms by which thalamocortical circuitry regulates the dynamic properties of cortical beta in PD are poorly understood. Using the recently developed generic dynamic causal modelling framework, we recursively optimised a set of plausible models of the thalamocortical circuit (n=144) to infer the neural mechanisms that best explain the transitions between low and high beta power states observed in recordings of field potentials made in the motor cortex of anesthetized Parkinsonian rats. Bayesian model comparison suggests that upregulation of cortical rhythmic activity in the beta-frequency band results from changes in the coupling strength both between and within the thalamus and motor cortex. Specifically, our model indicates that high levels of cortical beta synchrony are mainly achieved by a delayed (extrinsic) input from thalamic relay cells to deep pyramidal cells and a fast (intrinsic) input from middle pyramidal cells to superficial pyramidal cells. We therefore hypothesize that beta synchronisation at the cortical level could selectively be modulated via interventions that are capable of finely regulating cortical excitability in a spatial (delivered to either the superficial or deep cortical laminae) and time specific manner.

## Introduction

Neuronal oscillations are considered to be key elements of information flow (Buzsaki & Draguhn, 2004; Salinas & Sejnowski, 2001). For neural populations to communicate in a behaviour-specific and adaptive fashion, they may adapt their degree of rhythmic synchronization accordingly (Fries, 2005). In its normative physiological state, the Cortico-Basal Ganglia-Thalamo-Cortical circuit (CBGTC) exhibits transient (de-)synchronization in the beta band (13-30Hz) activity during motor control (Cassidy et al., 2002; Foffani et al., 2005; Pfurtscheller & Lopes Da Silva, 1999; Tsang et al., 2012; Zaepffel et al., 2013).

In Parkinson’s disease (PD), the loss of midbrain dopaminergic neurons in the substantia nigra pars compacta is thought to often promote excessive oscillatory synchronization of neuronal activity in the beta band across different nodes of the CBGTC circuit, which is thought to underpin some debilitating motor deficits such as bradykinesia and rigidity (Eusebio et al., 2009). When Parkinsonian motor deficits are attenuated with pharmacological (Levodopa) or neuromodulatory interventions (deep brain stimulation or optogenetics), a reduction in synchronization is observed in the beta-frequency band across different species, including humans (Brown, 2001; Eusebio et al., 2011; Kuhn et al., 2008; Levy et al., 2002; Priori et al., 2004; Silberstein et al., 2005), 1-methyl-4-phenyl-1,2,3,6-tetrahydropyridine treated non-human primate models of PD (Heimer et al., 2006; Nambu & Tachibana, 2014) and a 6-hydroxydopamine (6-OHDA)-lesioned rat model of PD (Gradinaru et al., 2009; Sharott et al., 2005).

Although excessive synchrony in the beta band (i.e. beta power) is traditionally described as a sustained event when averaged over seconds (Brittain & Brown, 2014; Brown, 2007; Lopez-Azcarate et al., 2010), it primarily manifests as intermittent events of high beta power or “beta bursts” (Feingold et al., 2015; Sherman et al., 2016; Tinkhauser et al., 2017; Little et al., 2012; Leventhal et al., 2012). Beta bursts have been defined operationally as epochs of beta oscillations that surpass a certain threshold – and their presence has been quantified in physiological (Sherman et al., 2016; Feingold et al., 2015) and pathological neural activity (Tinkhauser et al., 2017; Little et al., 2012). In Parkinson’s disease, the probability of long beta bursts has been positively correlated with PD motor symptom severity (Tinkhauser et al., 2017; Little et al., 2012).

Adaptive Deep Brain Stimulation (aDBS) is an intervention that has been developed to account for the transient nature of pathological neural synchrony in the beta band. In contrast to conventional DBS (cDBS), which continuously delivers high-frequency stimulation, aDBS adapts stimulation delivery according to the level of beta power (Little et al., 2013, 2016; Rosa et al., 2015), showing greater clinical efficiency (higher motor symptom relief and fewer secondary effects) than cDBS and random stimulation (Little et al., 2013).

From a neuronal network perspective, several studies have proposed that altered basal-ganglia output leads to excessive beta synchrony and motor impairments in PD (Bevan et al., 2002; Holgado et al., 2010; McCarthy et al., 2011; Terman et al., 2002). Employing Dynamic Causal Modelling (DCM) (Friston, 2003), a framework for specifying, fitting and comparing mathematical models of neural circuitry, Moran et al., 2011 and Marreiros et al., 2013 indicated modulation of the hyperdirect pathway and the projection from the subthalamic nucleus and globus pallidus externus as potential mechanisms for beta power enhancement in dopamine-depleted states.

Some experimental studies, on the other hand, support the role of cerebral cortex in the generation and modulation of beta oscillations (Jensen et al., 2005; Yamawaki et al.,2008). This perspective has motivated the consideration of cortical interlaminar interactions in the regulation of beta power. In the healthy state, the generation and modulation of beta oscillations has been investigated using DCM (Bhat et al.,2016), revealing a link between a set of laminar specific interactions within the primary motor cortex and the enhancement/suppression of beta power evoked by movement. Using a theoretical model, Sherman et al., 2016 suggested that high beta power events in the physiological state emerge through cortical laminar interactions conditioned by temporal characteristics of the distal and proximal synaptic drives in the neocortex.

Motivated by these studies, we hypothesized that – in the Parkinsonian state – an alteration of interlaminar and laminar-specific connectivity in the Thalamocortical (TC) loop contributes to the mechanisms generating the parkinsonian spectral profile. We focused on the TC loop due to the anatomical and functional characteristics of this network: 1) the cortex is an optimal target for non-invasive therapeutic techniques such as TMS and TACS (Barker et al., 1985; Cantello et al., 2002; Kobayashi & Pascual-Leone, 2003; Herrmann et al., 2013); 2) the thalamus is the only CBGTC node projecting directly to cortex, allowing for the integration of information from subcortical structures to the motor cortex (Wise & Donoghue, 1986; Brazhnik et al., 2016) and 3) cortex and thalamus establish a reciprocal relationship (Hooks et al., 2013), which is thought to play a key role in physiological and pathological sensory and motor computations (Sherman & Guillery, 2009). In PD, where motor impairments are the cardinal symptoms, understanding the synaptic dynamics and organization of the thalamocortical (TC) circuit could potentially shed light on pathophysiological mechanisms. Accordingly, we used cross spectral density (CSD) - DCM (Moran et al., 2009, 2011) with a neural mass model of the TC loop (Van Wijk et al., 2018) to characterize its contribution to spontaneous beta power fluctuations observed in the motor cortex of 6-OHDA-lesioned Parkinsonian rats.

## 2. METHODS

### 2.1. Electrophysiological recordings in Parkinsonian rats

The spectral data used in this study was based on motor cortex field potentials (electrocorticograms) recorded in 36 urethane-anesthetized rats rendered Parkinsonian by unilateral 6-OHDA lesions of midbrain dopaminergic neurons. To record electrocorticogram (ECoG) data, a steel screw electrode was implanted over the right somatosensory-motor cortex ipsilateral to the 6-OHDA lesion, and referenced to a steel screw electrode implanted over the ipsilateral cerebellar hemisphere. Electrophysiological recordings were carried out 21-42 days after surgery for the induction of 6-OHDA lesions, thus allowing for changes in the CBGTC circuit to stabilize. For detailed descriptions of electrode implantation, anaesthesia, surgical induction of 6-OHDA lesions and related procedures, please refer to (Mallet et al. 2008a, 2008b; Sharott et al., 2017). Only ECoG recordings made during periods of spontaneous ‘cortical activation’ were considered in this study (Mallet et al., 2008a, 2008b; Sharott et al., 2017). All experimental procedures were carried out on adult male Sprague-Dawley rats (Charles River, Margate, UK) and were conducted in accordance with the Animals (Scientific Procedures) Act, 1986 (UK).

### 2.2. Data processing

All operations described in this section were performed in Matlab. Recordings were down-sampled to 1000 Hz from 16000 Hz. To characterize the spontaneous beta power fluctuations typically observed in PD, we defined two conditions based on instantaneous beta power – condition one being Low Beta (LB) power and condition two being High Beta (HB) power. These conditions were based on fluctuations in beta power that enabled us to select data-features (i.e., timeseries) for subsequent dynamic causal modelling that were representative of the two conditions.

To extract the beta power envelope, we applied a second order band-pass Butterworth filter with cut-off frequencies at 15-35Hz to the ECoG recording and subsequently employed the Hilbert transform to compute the envelope of the ECoG in the beta frequency band. Each envelope was then divided into non-overlapping epochs of 500 msec. The two conditions were subsequently derived from the area under the envelope across the 500msec epochs: (1) LB epochs consisted of segments whose envelope area fell below the 5th percentile of the envelope area observed across all epochs, and (2) HB epochs consisted of segments whose envelope area was above the 95th percentile of the envelope area observed across all epochs (Fig.1). From each recording, we randomly selected 5 epochs per condition (n=5). This number corresponds to the minimum number of epochs found in either of the two conditions across all recordings.

**Figure. 1.**
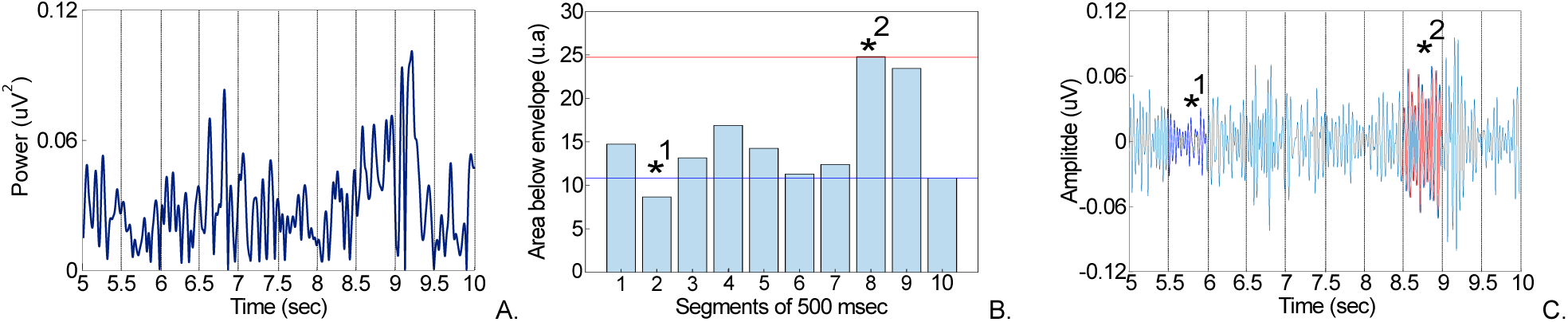
Extraction of low beta and high beta power features isolated from ECoG data. Panel A. shows the segmentation of the envelope into 500 msec epochs (5 seconds as an example). Panel B. depicts the area under the beta band envelope for each epoch. If an epoch was below the 5^th^ percentile of the area observed across all epochs (blue line), it was classified as low beta (*1); if an epoch was above the 95^th^ percentile of the area observed across all epochs (red line), it was classified as high beta (*2). Epochs in between the two percentiles were not considered. Panel C. shows the corresponding low beta (dark blue) and high beta (red) epochs in the ECoG signal filtered at 15-35 Hz.

### 2.3. Dynamic Causal Modelling (DCM)

DCM for cross spectral density is used to infer the hidden (neuronal) states (*z*) and synaptic parameters (*θ*) that generate spectral features of observed data (*u*) (Moran et al., 2009, 2011). Hidden states and unknown parameters cannot be observed directly but can be estimated under a generative or forward model. This model comprises a biophysical neural mass model and the spectral composition of neural and channel noise (Moran et al., 2008). The neural mass model is expressed in terms of a differential equation with the following form:

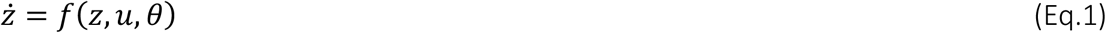

The neural mass model (*f*) – together with a likelihood model mapping hidden states to observed measurements – constitutes a generative model; namely, a probabilistic mapping between neural fluctuations and the spectral content of observed activity. Using a Bayesian framework, DCM estimates the (posterior) probability density over the synaptic parameters, which are the most likely value of the hidden parameters, given the observed data (Moran et al., 2011). The generative (neural mass) model calls on its biophysical parameters to describe the evolution of voltages (*v*) and currents (*i*) in each subpopulation of neurons (Jansen & Rit, 1995). In addition to estimating the posterior density over model parameters (e.g., synaptic connection strengths and the amplitude of neuronal fluctuations), DCM also provides an estimate of the evidence for a particular model or network architecture implicit in the generative model. This allows one to compare different models or hypotheses using Bayesian model comparison. A complete description of the mathematical framework that underwrites DCM can be found in (Moran et al., 2013).

#### 2.3.1. Neural mass model of the Thalamocortical circuit

A neural mass model of the Thalamocortical circuit was created comprising two formally distinct neural mass models of the motor cortex and the thalamus using the new generic framework for Dynamic Causal Modelling (van Wijk et al 2018) (Fig.2). Here, we adopted the motor cortex microcircuit (MMC) model developed by Bhat and colleagues (2016) and coupled it to a model of the thalamus, based on thalamic anatomical literature (Shepard & Grilner,2010; Douglas & Martin,2001). As with previous models of the sensory cortex – and incorporating the work of Yamawaki et al.,2014 - Bhat and colleagues (2016) used 3 excitatory subpopulations (neuronal ensembles consisting of “superficial”, “middle” and “deep” pyramidal cells located in the supragranular, granular and infragranular cortical layers, respectively) and one common inhibitory subpopulation (inhibitory interneurons) to model the primary motor cortex. In the MMC model, the coupling between these subpopulations (GABAergic or glutamatergic synapses) is tailored according to synaptic characteristics of the primary motor cortex: a reciprocal connection between superficial and middle pyramidal cells (Yamawaki et al., 2014), a reciprocal connection between superficial and deep pyramidal cells (Hooks et al., 2013; Yamawaki & Shepherd, 2015; Anderson et al., 2010, Weiler et al., 2008), a reciprocal connection between each of the three pyramidal subpopulations and the inhibitory subpopulation (Fino et al., 2013), and a cell type specific self-inhibitory connection (Bastos, 2012; Yoshimura & Callaway, 2005). In this study, the thalamus was modelled using an excitatory subpopulation (neuronal group of thalamic relay cells) and an inhibitory subpopulation (neuronal group of thalamic reticular cells) (Shepherd & Grillner, 2010) that were connected as follows: a reciprocal connection between relay and reticular cells (Harris, 1987; Cox et al., 1997) and a self-inhibitory connection of reticular cells (Shu & McCormick, 2002). Although we acknowledge that there are distinct thalamic nuclei (i.e. neuronal ensembles receiving afferents from different brain regions (Sherman & Guillery, 2001) the thalamus was modeled here as a single neuronal mass model. An important extension of the current work would be to subdivide the motor thalamus (ventral anterior, ventral lateral and ventral medial nuclei in rodents) into input zones that receive GABAergic drive from the Basal-Ganglia and glutamatergic drive from the cerebellum (Kuramoto et al., 2011; Nakamura et al., 2014).

**Figure. 2.**
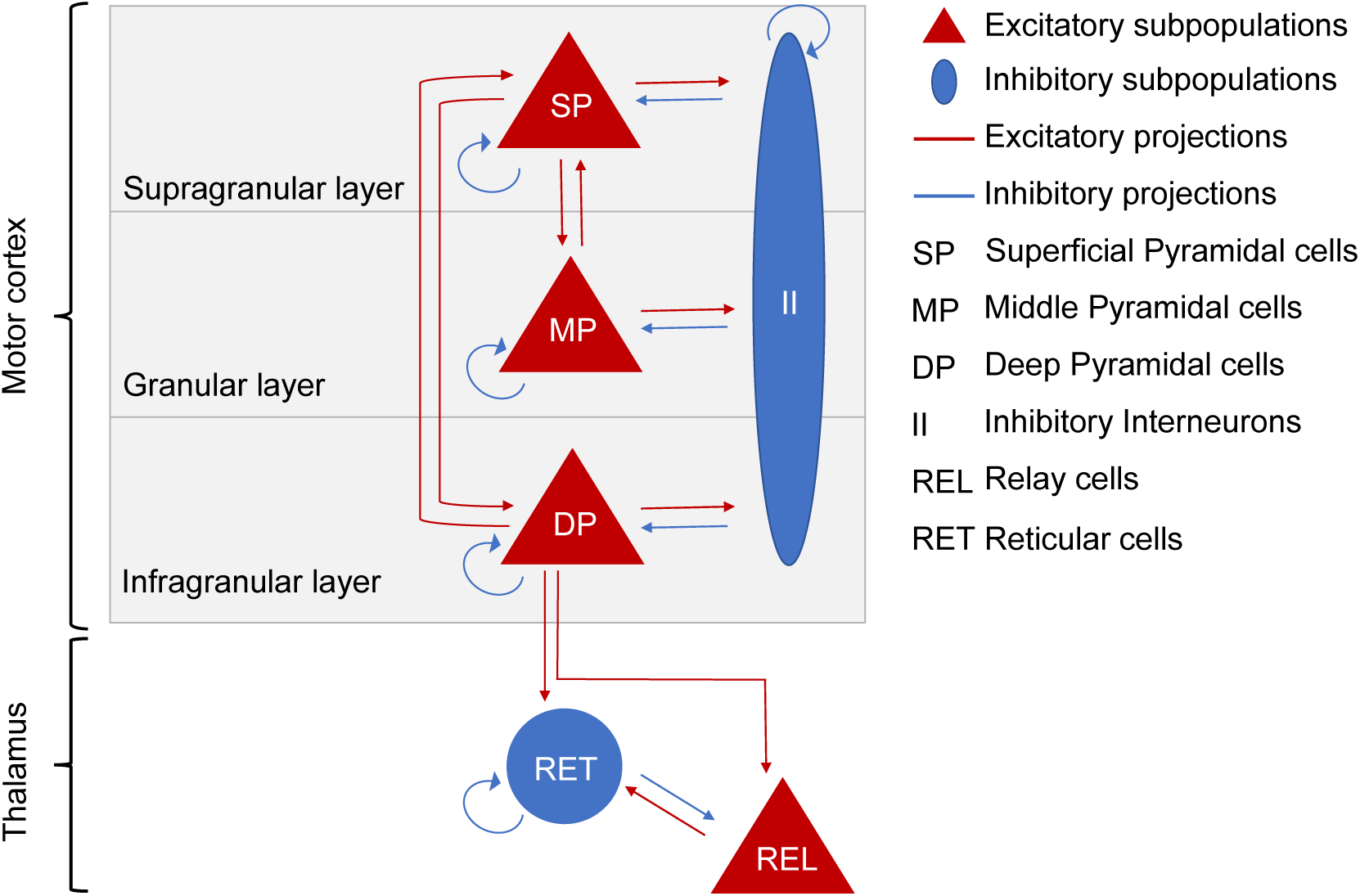
Sources, subpopulations and synaptic projections of a thalamocortical loop neural mass model. The top part of the diagram describes the first source - motor cortex and its subpopulations: superficial pyramidal cells (SP) in the supragranular layer, middle pyramidal cells (MP) in the granular layer, deep pyramidal cells (DP) in the infragranular layer and, inhibitory interneurons (II) as a common inhibitory subpopulation to the 3 cortical laminae. Intrinsic synaptic connections among the above subpopulations comprise a reciprocal connection between superficial and middle pyramidal cells, a reciprocal connection between superficial and deep pyramidal cells, a reciprocal connection between each of the three pyramidal subpopulations and the inhibitory subpopulation and finally, a self-inhibitory connection to each cortical node. The bottom part of the diagram depicts the thalamus and its subpopulations: reticular thalamic cells (RET) as the inhibitory subpopulation of the thalamus and relay cells (REL) as the excitatory subpopulation of the motor thalamus. Intrinsic synaptic connectivity of the thalamus comprises a reciprocal connection between relay and reticular cells and self-inhibitory connection of reticular cells. As corticothalamic extrinsic connections, deep pyramidal cells were considered to send afferents to both relay and reticular subpopulations, while the model space for thalamocortical projections is described in section 2.3.4 and illustrated in Fig.3 (top panel).

To model the extrinsic synaptic interactions between the motor cortex and thalamus we used two corticothalamic projections from deep pyramidal cells to thalamic relay cells and thalamic reticular cells (Bourassa et al., 1995; Jones, 2001). Although the thalamus is thought to project to all layers of the cortex (Hooks et al., 2013) and the ventromedial nucleus (VM) of the motor thalamus has been shown via immunochemistry studies to project mainly to layers I and II of the motor and anterior cingulate cortices (Arbuthnott et al., 1990; Clascá et al., 2012; Kuramoto et al., 2015), it is not clear which thalamocortical projections are important in modulating beta power. To resolve this, we considered different models to test the impact of including different connections on model evidence (section 2.3.4.).

#### 2.3.2. Neural dynamics

In DCM, neural dynamics (i.e., fluctuations in voltages and currents) at the subpopulation level is described by two key operations (Eq.2): a convolution operator and an output operator (Moran et al., 2007). The convolution operator transforms presynaptic inputs (firing rate) into postsynaptic membrane potentials based on a synaptic impulse response function, which considers the nature of the synapse (i.e. excitatory or inhibitory).

The output operator consists of a non-linear function that converts the postsynaptic membrane potentials into a firing rate to be relayed to another subpopulation. This is conveyed through a sigmoid function *S* which captures the membrane sensitivity and firing threshold of each subpopulation. Furthermore, the shape of the sigmoid function (slope) measures the efficacy of a presynaptic ensemble to generate output. This output is additionally scaled by the synaptic coupling strength as illustrated by the following generic second order differential equation:

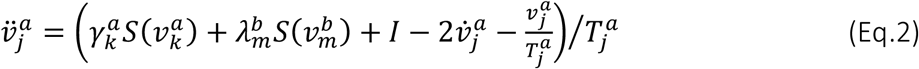

Here, the averaged membrane potential *v* of the subpopulation *j* in the source *a* is influenced by subpopulations of the same source with synaptic strength *γ* and subpopulations from different sources with synaptic strength *λ*. Intrinsic synapses *γ* show a positive synaptic strength if glutamatergic and negative synaptic strength if GABAergic. *S* denotes the sigmoid function above and *T* the subpopulation-specific membrane time constant. Endogenous fluctuations or input, *I* is modelled as a mixture of white and pink noise and drives middle pyramidal cells and thalamic relay cells.

In this study, we used DCM for cross spectral density (Moran et al., 2009, 2011) where the data generated by a model of neural hidden states are expressed as cross spectra in channel space (ECoG screw electrodes). The mapping between the hidden neural states and measured signals is based on an observation function, where a gain matrix *L* scales different contributions (here [0.6 0.2 0.2]) from superficial, middle and deep pyramidal cells respectively to generate the observed ECoG signal. A summary of these parameters and their prior values are shown in Table.1. Prior values were based on previous DCM studies (Bhat et al. 2016) and optimized for our study.

**Tabel 1.**
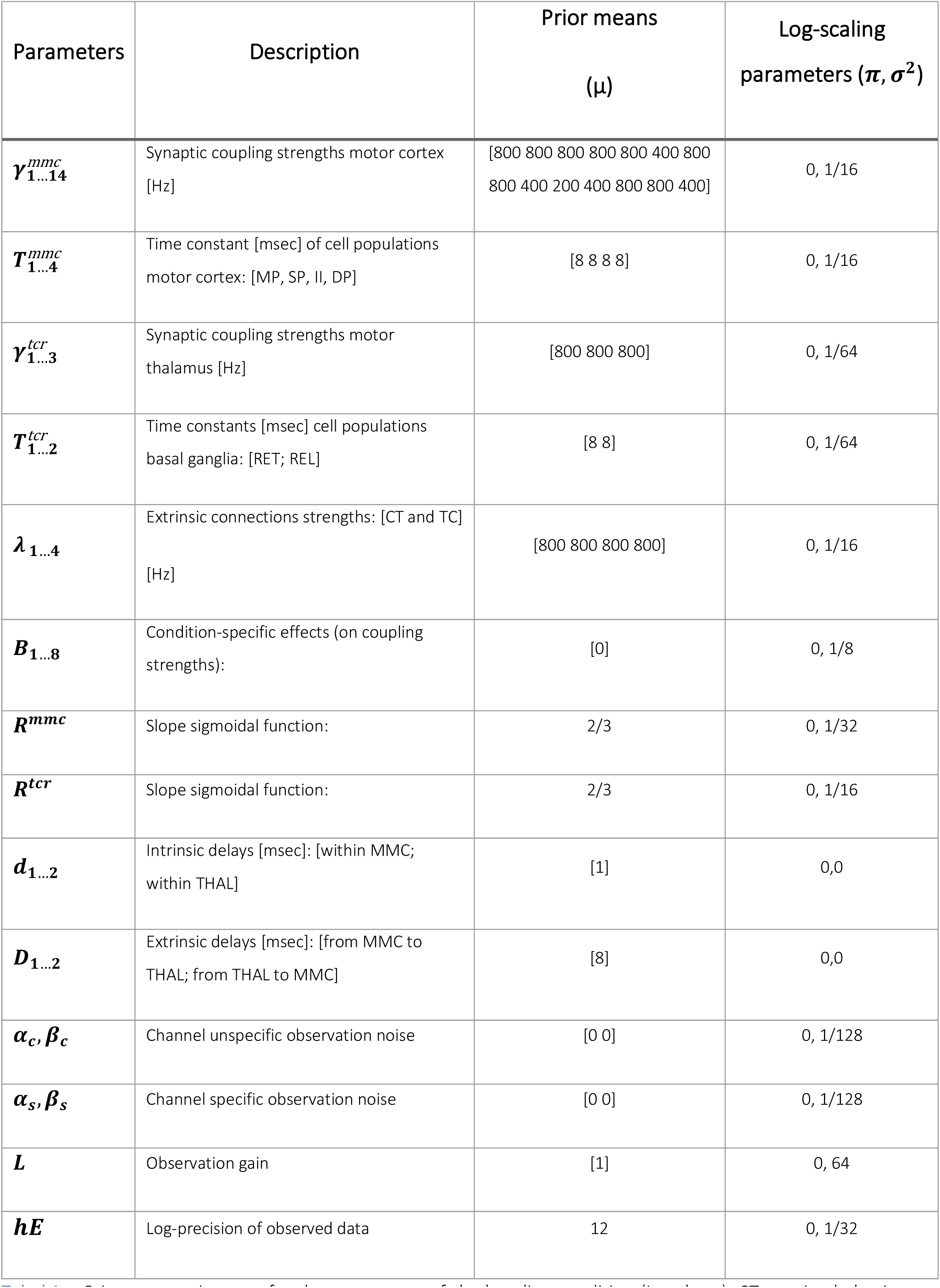
Prior expectations set for the parameters of the baseline condition (Low beta). CT-corticothalamic projections; TC-thalamocortical projections; MMC – motor microcircuit; THAL-thalamus.

#### 2.3.3. Model inversion

In DCM, model inversion iteratively tunes the model’s parameters to optimize the fit of the predicted electrophysiological data to the observed data. Using a standard (variational) Bayesian scheme, model inversion uses priors to constrain the search of parameter space to explain the observed spectral features of electrophysiological data. When fitting the data (i.e., inverting the model), the optimization of model parameters uses a variational Laplace scheme to minimize a (free energy) bound on (negative) log model evidence. This free energy approximation to model evidence is subsequently used for model comparison (Friston et al., 2007; Friston & Stephan, 2007).

In brief, model evidence is the (marginal) likelihood of observing data given a model, E(F|H). It reflects a balance between accuracy (goodness of fit between predicted and observed spectral densities) and complexity (divergence between prior and posterior parameter estimates) (Stephan et al., 2010). This balance depends upon the expected precision of the observed data. Given the high signal to noise ratio in the data obtained using the electrocorticographic recording method, the expected precision of observed data was assumed to be high (with a log precision of 12).

At this stage, low beta power was set as our baseline condition (with prior expectations optimized to best explain its spectral features). Condition-specific effects (*B parameters*) on both extrinsic (between-regions) and intrinsic (within-regions) coupling strengths (Moran et al., 2007) were used to explain periods of high beta power. In other words, we estimated the changes in synaptic efficacy required to move from a low beta power condition to a high beta power condition.

#### 2.3.4. Bayesian Model Comparison and parameters analysis

A set of models were implemented which varied according to 2 factors: i) the laminar-specificity of thalamocortical projections that generate beta oscillations, and ii) the changes in synaptic connectivity within the TC loop (intrinsic and/or extrinsic) required to induce a transition from a low beta power condition to a high beta power condition.

The first factor comprised 9 families (types) of models. These models had identical intrinsic and corticothalamic connections as described in section 2.3.1 and illustrated in (Fig.2) but differed in the laminar targets of thalamocortical afferents: 1) superficial pyramidal cells; 2) middle pyramidal cells; 3) deep pyramidal cells; 4) superficial plus middle pyramidal cells; 5) middle plus deep pyramidal cells; 6) superficial plus deep pyramidal cells; 7) superficial pyramidal cells plus inhibitory interneurons; 8) middle pyramidal cells plus inhibitory interneurons and 9) deep pyramidal cells plus inhibitory interneurons (Fig.3, top panel).

**Figure. 3.**
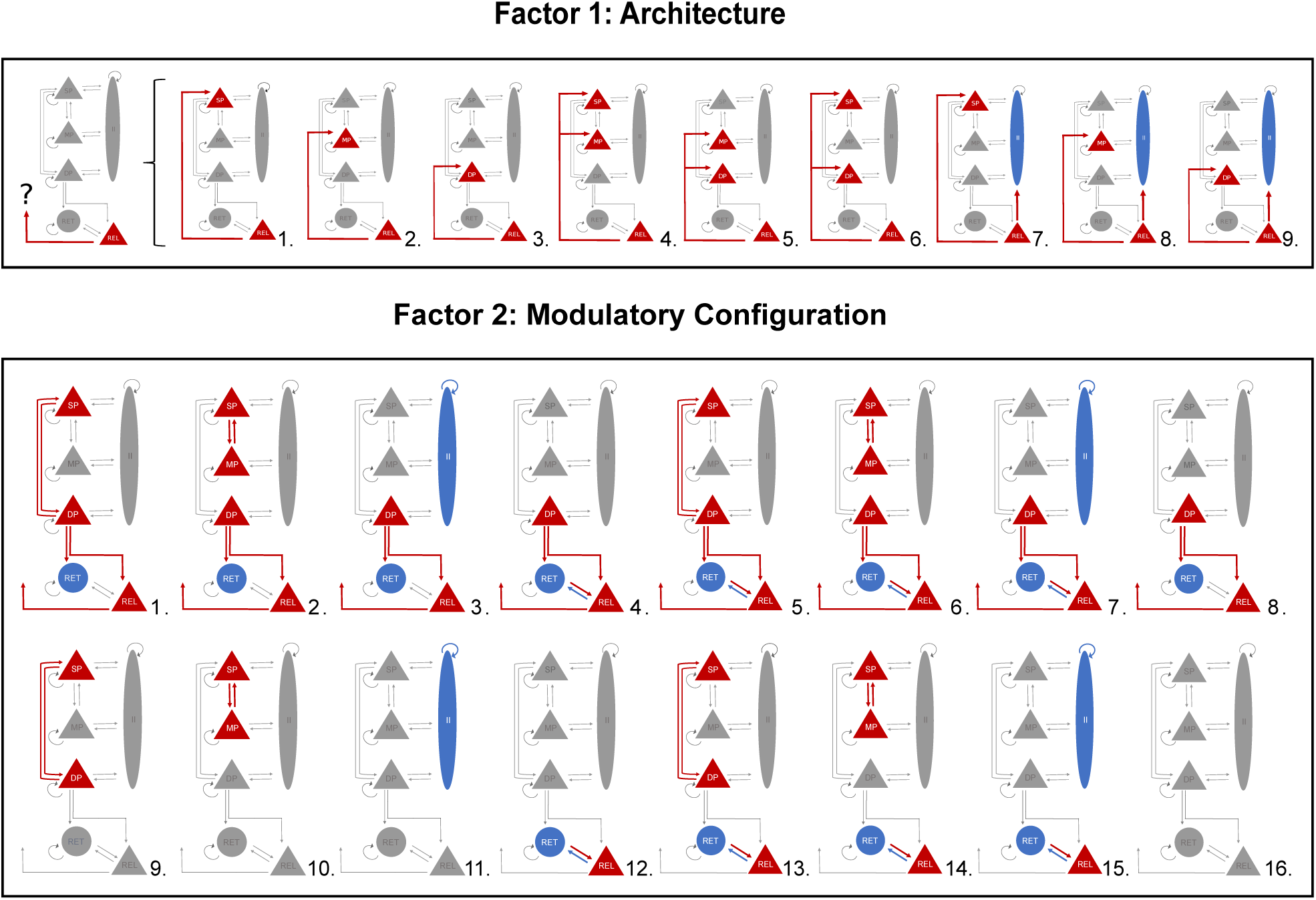
Competing models of the Thalamocortical circuit as described by factors 1 and 2 (9 architectures times 16 modulatory configurations). The diagram on the top (factor 1: architecture) describes the 9 families of models constructed to elucidate which thalamocortical projections are the most plausible explanation for the generation of beta oscillations (1.-3.) accounts for a singular projection from thalamus to motor cortex via superficial pyramidal cells, middle pyramidal cells and deep pyramidal cells; (4.-6.) accounts for two afferents to two excitatory subpopulations of the motor cortex via superficial and middle pyramidal cells, middle and deep pyramidal cells and superficial plus deep pyramidal cells, and (7.-9.) accounts for projections to the superficial pyramidal subpopulation and inhibitory interneurons, the middle pyramidal subpopulation and inhibitory interneurons and deep pyramidal cells and inhibitory interneurons. To disclose the synaptic modulation (intrinsic and/or extrinsic) responsible for an enhancement of beta power, the models on the bottom (factor 2: modulatory configuration) feature 16 different modulatory configurations, under each of the 9 architectures described above. The first eight set of connections (1.-8.) entail extrinsic and intrinsic synaptic modulation (except for model 8, with no intrinsic modulation) and the second eight set of connections (9.-16.) considers intrinsic modulation only. The intrinsic modulatory connections in family 2 were: (1. and 9.) cortical modulation via reciprocal connection between superficial and deep pyramidal subpopulations; (2. and 10.) cortical modulation via reciprocal connection between superficial and middle pyramidal subpopulations; (3. and 11.) cortical modulation via self-inhibitory connection of the inhibitory interneurons subpopulation; (4. and 12.) thalamic modulation via reciprocal connection between reticular cells and relay cells; (5. and 13.) cortical and thalamic modulation via reciprocal connection between superficial and deep pyramidal subpopulations plus reciprocal connection between reticular cells and relay cells; (6. and 14.) cortical modulation via reciprocal connection between superficial and middle pyramidal subpopulations plus reciprocal connection between reticular cells and relay cells; (7. and 8.) self-inhibitory connection of the inhibitory interneuron subpopulation plus reciprocal connection between reticular cells and relay cells and lastly, (8. and 16.) the null hypothesis that neither extrinsic nor intrinsic connections change to explain condition specific changes in cortical beta power (i.e. enhancement of beta).

The second factor comprised 16 families of models that varied in the set of connections that could show condition specific effects (i.e. B matrix). For each one of the 9 architectures in the first factor, we explored condition specific effects by including or not the following features: intracortical modulatory synapses; intrathalamic modulatory synapses and extrinsic (between cortex and thalamus) modulatory synapses (Fig.3, bottom panel). There were therefore 9 x 16=144 candidate models in total.

Bayesian Model Comparison (BMC) was used to determine the model with the highest log-model evidence among the models described above (Stephan et al., 2010). We then characterised the parameters of the winning model at the group level using Parametric Empirical Bayes (PEB) (Friston et al.,2015). Here, only a subset of parameters was taken to the group level and assumed to exhibit random effects: synaptic coupling strength of intrinsic and extrinsic connections (G and A parameters in the DCM respectively); condition-specific effects on coupling strength (B parameters) and the time constants of subpopulations (T).

## 3. RESULTS

### 3.1. Model Selection

The model with the highest evidence for the transition from low beta epochs to high beta epochs (Fig.4) was that with i) an architecture featuring thalamocortical projections from the thalamic relay subpopulation to both deep pyramidal and inhibitory interneuron subpopulations in cortex and ii) modulatory changes in: intrinsic connections at the cortical level between superficial and middle pyramidal cells; intrinsic connections at the thalamic level, between relay and reticular cells; corticothalamic extrinsic connections from deep pyramidal cells to relay and reticular cells and thalamocortical connections from relay cells to deep pyramidal and inhibitory interneurons. The winning model shows a free energy difference (i.e., log Bayes factor) of approximately 6 from the next closest model (Fig.S.2). This corresponds to very high evidence for the winning model, in relation to alternative explanations.

**Figure. 4.**
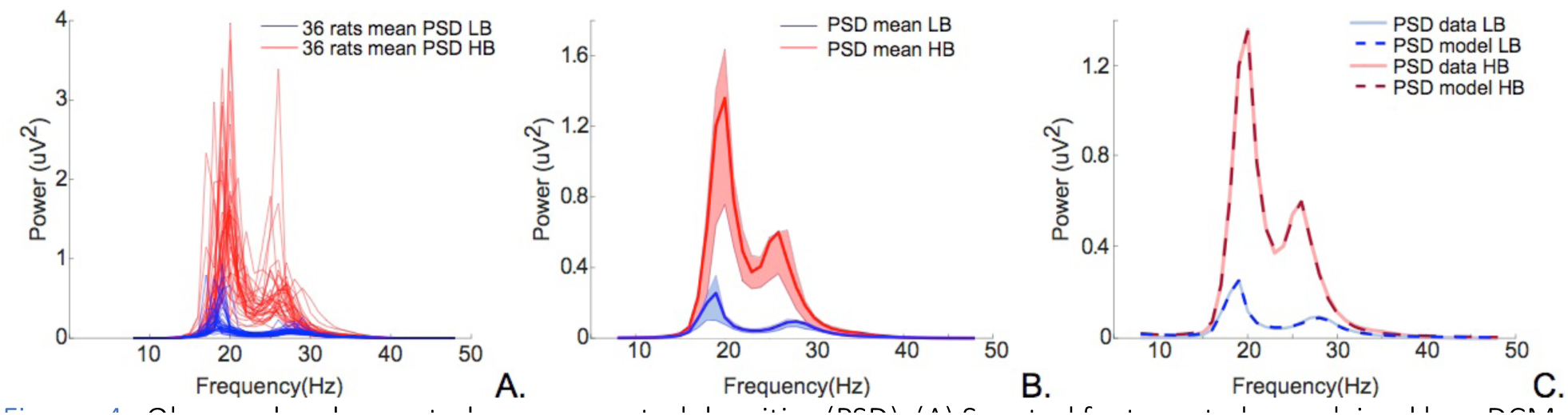
Observed and expected power spectral densities (PSD). (A) Spectral features to be explained by a DCM: Red lines depict the mean high beta spectral densities and blue lines the mean low beta spectral densities from each of the 36 rats. (B) Group mean of HB spectral densities in red and LB spectral densities in blue. Respective variabilities (75^th^ and 25^th^ percentiles of the mean spectra) denoted in light red and light blue. (C) Goodness of the fits between mean data spectral densities and spectral densities generated by the winning model. The full red line shows the mean of high beta data and the dark red dashed line the high beta spectra estimated by the winning model (correlation coefficient, r=0.9997). The full dark blue line refers to the mean of low beta data and the dark blue dashed line to the low beta spectra produced by the winning model (correlation coefficient, r=0.9957).

Using fixed-effects Bayesian Model Comparison (FFX-BMC) to make inferences at the family level, the architecture with thalamic projections to DP and II showed the highest evidence across subjects (Fig.5.A). Similarly, the condition specific effects in the reciprocal connection between SP-MP, REL-RET and DP-REL plus connections from DP-RET and REL-II had the highest posterior probability (Fig.5.B). These results confirm our hypothesis that both the laminar-specificity of extrinsic connectivity and intrinsic connections are key elements underlying the modulation of oscillatory activity in the beta band.

**Figure. 5.**
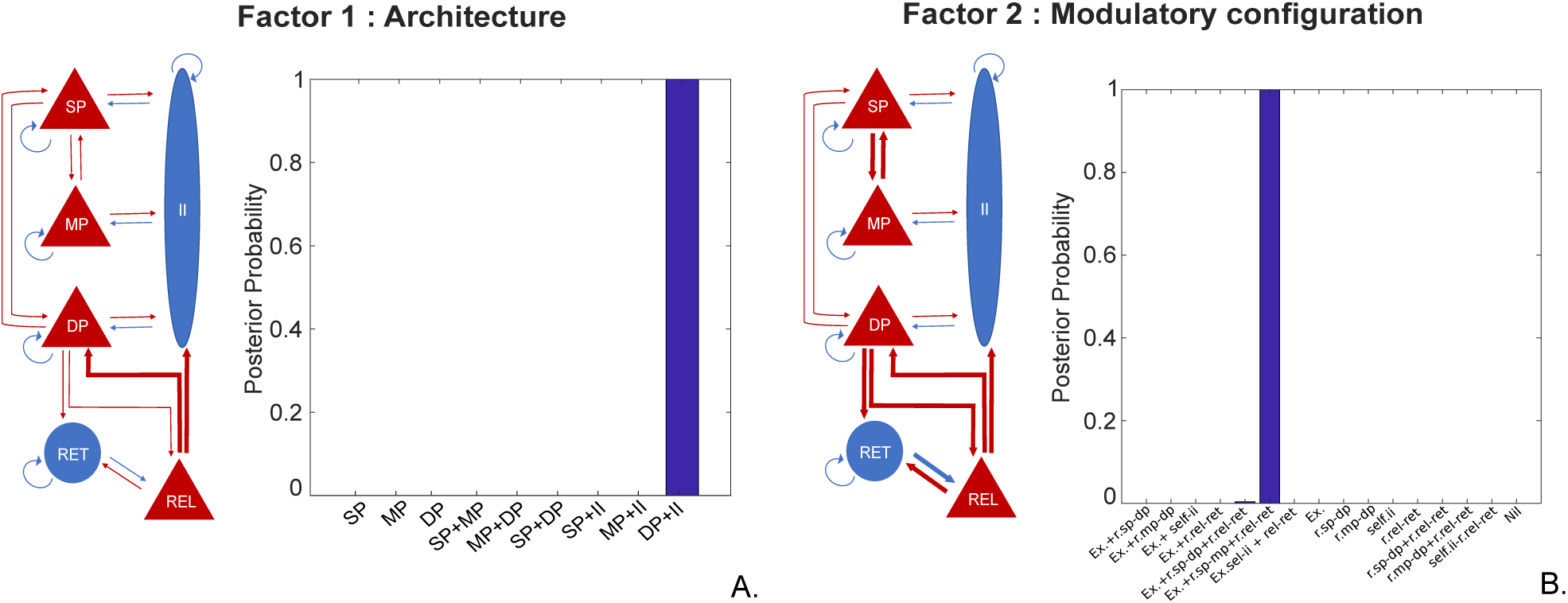
Schematic and posterior probability of the winning model selected via FFX-BMC. Diagram and bar plot (A) refer to architecture of the winning model. These results suggest that thalamocortical projection to the deep pyramidal cells and cortical inhibitory subpopulation (in thick lines) were crucial for the generation of beta oscillations and that this effect was consistently observed across subjects (posterior probability of 1). Diagram and bar plot (B) indicate the modulatory connections of our winning model (BMC factor two). The diagram shows the set of connections as thick lines to have a higher likelihood (compared to the homologous 15) of inducing the power spectral changes observed (beta enhancement). These being: a reciprocal connection between superficial and middle pyramidal subpopulations, reciprocal connection between thalamic relay and reticular subpopulation and a reciprocal extrinsic connection between deep pyramidal cells and thalamic relay cells, an extrinsic connection from deep pyramidal cells to thalamic reciprocal cells and from thalamic relay cells to cortical inhibitory interneurons. The bar plot shows a posterior probability greater than 0.99 for the modulatory configuration described above and a negligible posterior probability of approximately 0.004 for a modulatory configuration which assumed the same modulatory characteristics as the winning model except for the intrinsic synaptic mechanisms of the motor cortex; i.e., presenting an intracortical modulation via reciprocal connections between superficial and deep pyramidal cells instead of reciprocal connections between superficial and middle pyramidal cells. (Ex.-extrinsic connections, r.-reciprocal connections, sp.-superficial pyramidal cells, mp.-middle pyramidal cells, dp.-deep pyramidal cells, ii.-inhibitory interneurons, rel. – relay cells and ret. – reticular cells).

### 3.2. Parameter analysis

The results from our second level analysis (PEB modelling of A,G,B and T parameters at the group level) suggest that the transition from low beta state to high beta state is induced by i) an increase in synaptic strength in connections from relay cells to inhibitory interneurons, relay cells to deep pyramidal cells, middle pyramidal cells to superficial pyramidal cells and relay to reticular cells; plus ii) a reduction of synaptic strength in connections from superficial to middle pyramidal cells, deep pyramidal to both relay and reticular cells and from reticular to relay cells (Fig.6). Additionally, from the posterior distribution of our B parameters we assessed the effect size of each modulatory connection on the enhancement of beta – illustrated in the bar plot below (Fig.6).

**Figure. 6.**
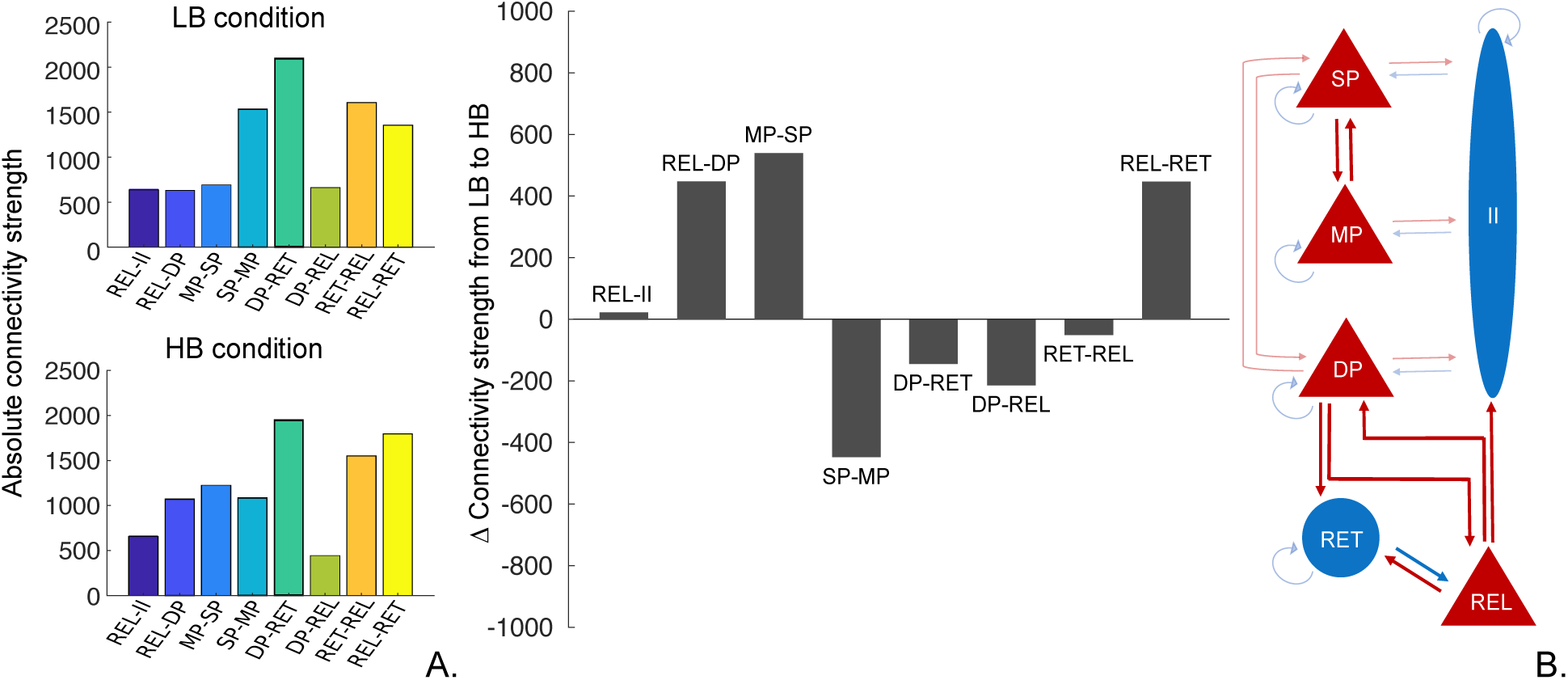
Average modulatory effect of B parameters (condition-specific parameters) obtained via Parametric empirical Bayes analysis (Friston et al.,2015). The two bar plots on the left-hand side illustrate the absolute connection strength of each modulatory connection in the low and high beta conditions. The bar plot in the centre shows how connectivity strength of B parameter changed at the group level in order to induce an increase of beta power. Negative values of change indicate a reduction in connectivity strength and positive values an increase. The anatomy of these connections is illustrated in the diagram on the right-hand side.

Considering the connections that showed the greatest change to explain beta enhancement: REL-DP, MP-SP and REL-RET; we further analyzed, via forward modelling, the impact of simultaneous alteration of the above connection strengths on the magnitude of beta power. As such, we aimed to characterize the contribution of these three parameters to the gradual transition between the two states – low and high beta power. This *post hoc* simulation allows us to characterize the selective effect of specific connections on the expression of cortical beta power level. Fig.7.A, suggests that when the intrinsic thalamic connection from relay to reticular cells have low levels of coupling strength, high beta power appears abruptly as the coupling strength from middle to superficial pyramidal cells is increased. This emergence of beta is modulated by the gradual change in coupling strength from thalamic relay cells to deep pyramidal cells, where reduced coupling leads to higher levels of beta.

**Figure. 7.**
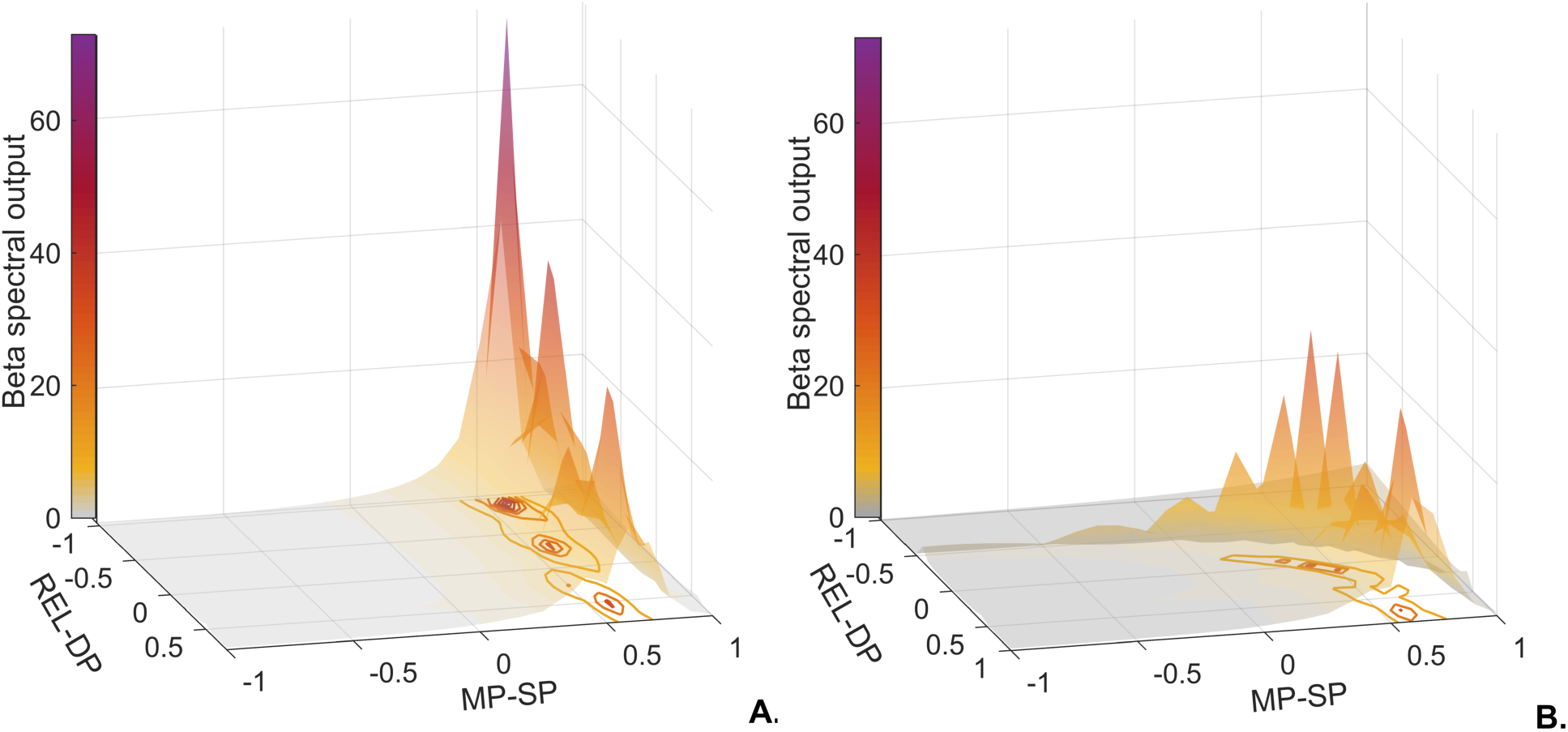
Exploration of parameters space of connections with the greatest effect on beta enhancement. Plot A. shows the impact of changing the coupling strength of MP-SP and REL-DP on the beta spectral output, when the coupling between *REL-RET is weak*. Here, although an increase in synaptic strength from MP-SP is enough to generate relatively high levels of beta spectral output, the connection from REL-DP seems to have a modulatory effect, i.e., the weaker the extrinsic coupling between relay and deep pyramidal cells the higher the beta spectral output. Plot B. considers a (constantly) strong coupling between *REL-RET* with the same changes in coupling strength between MP-SP and REL-DP. This time, we observe that an increase in beta is achieved with a concurrent strengthening of both MP-SP and REL-DP connections. In both plots, axis x and y denote a reduced connectivity strength when values are between −1 and 0 and an increased connectivity strength when values are between 0 and 1 for connections from middle pyramidal cells to superficial pyramidal cells and from relay cells to deep pyramidal cells respectively. Axis Z and colormap depict the magnitude of beta power.

On the other hand, Fig.7.B, suggests that when the connectivity from relay to reticular cells is high, a concurrent increase in the coupling from both relay to deep pyramidal cells and middle to superficial pyramidal cells is required to achieve beta power enhancement. In short, the level of cortical beta depends on the increase in both superficial and deep pyramidal excitation.

We additionally explored the spectral output of each subpopulation of our TC neural mass model (Fig. 8) and observed that all nodes generated spectral curves within the beta band during both conditions and increased their power from condition one (LB) to two (HB) as expected. In particular, the populations yielding the largest beta enhancement in the motor cortex and thalamus were the deep pyramidal cells and relay cells respectively.

**Figure. 8.**
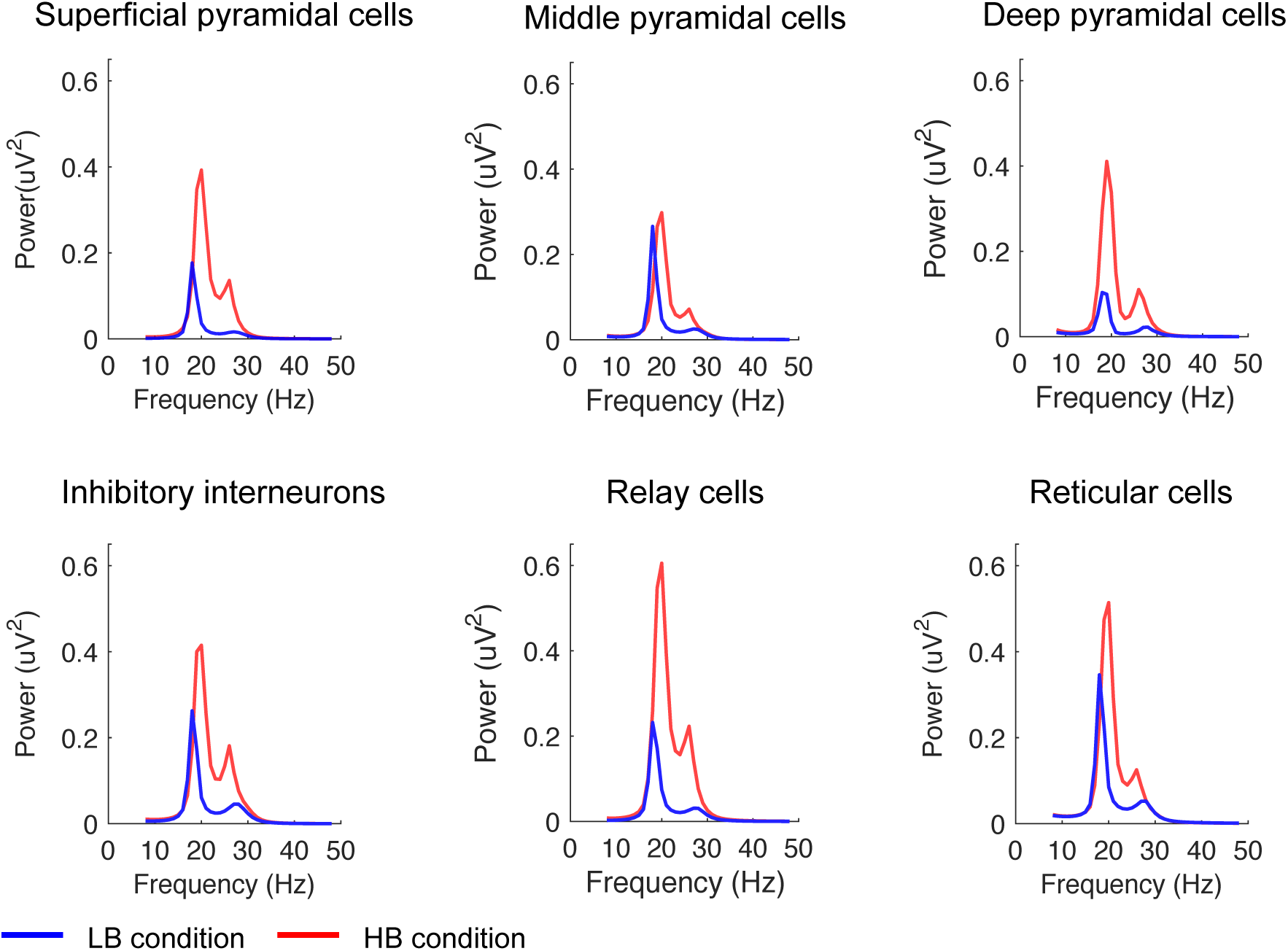
Spectral output of subpopulations at Low Beta (LB) and High Beta (HB). All neural groups (in cortex and thalamus) have generated spectral responses within the beta band in both conditions as expected. Together with an increase in power a small increase in frequency peak of approximately 1-2 Hz is also apparent – when comparing the output generated in condition 1 (LB) with condition 2 (HB). Note that deep pyramidal population of the cortex and relay population of the thalamus are the populations generating a relatively large beta enhancement.

## 4. Discussion

In this study, we aimed to identify the network mechanisms that contribute to the dynamic regulation of beta synchrony in the parkinsonian motor cortex. In-vivo studies of the basal ganglia thalamocortical (BGTC) circuit suggest that alterations in the firing rate across the direct and indirect pathways are responsible for the motor impairments observed in PD (Nambu, 2004; Smith al., 1998). Similarly, in-silico simulations of the BGTC circuit propose that an altered coupling from the subthalamic nucleus to globus pallidus externus, and strengthening of the hyperdirect pathway play an important role in the enhancement of beta synchrony following chronic dopamine depletion (Marreiros et al., 2013; Moran et al., 2011).

Our study complements the literature on PD, while exploring two novel concepts: i) laminar-specific dynamics within the motor circuit as a putative mechanism for the spontaneous modulation of beta power and ii) short-term synaptic processes, i.e. transient alterations in effective connectivity to be responsible for the spontaneous and intermittent nature of beta power observed in Parkinsonian time-series.

Focusing on the Thalamocortical loop of the BGTC circuit, we have employed DCM to identify a model of TC interactions that offers plausible substrates for the transient enhancement of cortical beta. Our study suggests two core features of the thalamocortical circuit that may underwrite the genesis of beta oscillations in the parkinsonian state: 1) laminar specific thalamocortical projections; and 2) modulation of synaptic strength across all network levels (i.e. within and between structures).

To model different levels of beta synchrony, we extracted low beta epochs and high beta power epochs from motor cortex ECoG recordings acquired from anesthetized rodents rendered Parkinsonian by 6-OHDA lesions. This rodent model is useful in capturing the chronic dopamine depletion that is common to ‘late stage’ PD, and has been widely used for studies of the mechanisms by which excessive beta synchrony arises and propagates within the BGTC circuit in Parkinsonism. Moreover, the abnormal beta oscillations present in the BGTC circuit in anesthetized and behaving 6-OHDA lesioned are similar in many respects to those present in unmedicated people with PD (Sharott et al, 2005; Mallet et al. 2008a, 2008b; Avila et al. 2010; Degos et al. 2009; Nevado-Holgado et al. 2014; Brazhnik et al. 2016; Sharott et al. 2017). We used DCM in this study since it quantifies effective connectivity changes underlying transient modulations in cortical beta power. Previously, Bhat et al., 2016 used DCM to link interlaminar dynamics within the motor cortex to the modulation of beta activity, evoked by movement. Fitting MEG data from healthy subjects to a neural mass model of the motor cortex, Bhat and colleagues reported that the increase in beta power observed due to the transition from grip to rest was induced by an increase in the extrinsic input applied to deep and superficial layers of the cortex. Our study suggests that beta power enhancement in Parkinsonism can be attributed to an increase in excitatory inputs to SP and DP; specifically from MP and thalamic relay cells, respectively – and a concomitant reduction of excitatory input to MP. In addition, our results highlight the importance of intrinsic interactions in the thalamus for beta power modulation as the excitatory projection from the thalamic relay cells to reticular cells also contributes to cortical beta enhancement (Fig.6).

Similarly, focusing on cortical intrinsic dynamics, Sherman et al., 2016 used a computational model to generate transient high beta power events (i.e. beta bursts), which were temporally identical to those observed in the somatosensory and frontal cortices in the physiological state. Two circuit features have been proposed as crucial for the generation of beta bursts: 1) a drive from the lemniscal thalamus to the proximal dendrites of the pyramidal neurons and inhibitory interneurons in L2/3 and L5 (via the granular layer) and 2) a strong drive from the nonlemniscal thalamus to the distal dendrites of the pyramidal neurons and inhibitory interneurons found in supragranular and infragranular layers.

It should be noted that due to the nature of the neural mass models employed in this study, we were not able to model detailed dendritic dynamics that contribute to the generation and modulation of neural activity in the beta band. Instead, here we assumed fixed conduction delays for all within-region connections (1 ms) and between-regions projections (8 ms), and did not account for variable propagation delays for inputs arriving to distal and proximal dendrites. This creates a distinction between the excitatory input received by the superficial pyramidal cells versus that received by the deep pyramidal cells; since the latter is attributed to an extrinsic projection from thalamus and hence is inherently modelled with longer conduction delays. Nonetheless, our results relate to the observations made in Sherman et al., 2016, given that comparable circuitry mechanisms yielded similar oscillatory effects. In other words, both studies propose that laminar specific excitation of the motor cortex must occur with two temporally separate inputs in order to achieve high beta power oscillatory activity.

Furthermore, it is worth noting that alternative models of the TC circuit (Fig.3) showed lower model evidence. Specifically, models, allowing for projections from the thalamic relay cells to both the superficial and deep layers of the motor cortex, had lower model evidences than the winning model. Possibly, this was because simultaneous projection from the thalamic relay cells to the superficial and deep layers would not have allowed for a differentiation in input delays contrary to the winning model. Detailed analysis of the parameter space on gradual beta increase, opposed to a transition from extremely low to extremely high beta power, further corroborated that the intrinsic (shorter delay) input from middle to superficial cells should have high levels of synaptic strength – together with the extrinsic (longer delay) input from the thalamic relay to deep pyramidal cells – to explain up-regulation of beta synchrony when the coupling from thalamic relay to reticular is strong. (Fig.7B). Nevertheless, taking both scenarios into account – strong vs weak connectivity from relay cells to reticular cells-excitability of superficial pyramidal cells via middle pyramidal cells must assume a moderate coupling strength to avoid a ramping up of beta synchrony at the cortical level; highlighting a potential substrate that could be targeted in order to control and modulate cortical beta.

An important difference between Sherman et al., 2016 and our study stems from the assumptions made on thalamic activity patterns. Sherman et al., 2016 posit that thalamic activity should be in the alpha band to drive beta bursts in the somatosensory and frontal cortices. However, in our study, thalamic neurons exhibited activity in the beta band during both low and high beta power conditions (Fig.8). Our results are supported by recent experimental work showing a substantial and coherent enhancement of beta activity (30-36Hz) in the motor thalamus and motor cortex of behaving 6-OHDA-lesioned rats (Brazhnik et al., 2016). There is also evidence of aberrant beta synchrony in the thalamus of unmedicated PD patients (Kempf et al. 2009) Taken together, these results emphasise that thalamic neural activity in the beta band is likely to be a contributing circuit feature for the generation of aberrant beta synchronization in PD, and highlight a functional coupling between the thalamus and deep layers of the motor cortex.

Our results also support the observation that deep pyramidal cells play a prominent role in the emergence of beta synchrony; since the deep pyramidal subpopulation exhibited a relatively large increase in beta spectral output between low and high beta power conditions (Fig.8). This observation is in line with a high-resolution MEG study by Bonaiuto et al.,2017, where a mapping between neural activity and cortical laminae was proposed. In this study, the modulation of beta activity during a motor task has shown a stronger signal component in the deep layers of the contralateral sensorimotor cortex than the superficial ones (Bonaiuto et al., 2017).

From a clinical perspective, our study provides insights into potential therapeutic strategies that could be utilized to modulate the network mechanisms responsible for the enhancement of cortical beta in PD. Specifically, we speculate that cortical stimulation aimed to reduce the excitability levels of either the superficial or deep pyramidal cells could be a potential non-invasive therapeutic strategy for PD.

## 5. CONCLUSION

A broadly accepted postulate concerning healthy and Parkinsonian states of the CBGTC circuit is that they exhibit differential patterns of synchronization at beta frequencies. (Brittain et al., 2014; Gatev et al., 2006; Hammond et al., 2007). While exaggerated beta synchronization has been associated with more frequent high power beta bursts in PD (Tinkhauser et al., 2017; Little et al., 2013), healthy states seem to manifest as an adequate balance between high and low beta bursts and therefore a flexible motor behaviour (Feingold et al., 2015; Sherman et al., 2016). Following this reasoning, a recognition of the mechanisms adopted by the CBGTC network to regulate beta spectral undulations is vital to better understand healthy and diseased states; and consequently, inform novel therapeutic strategies. Here, using DCM, we highlight a set of synaptic alterations in the thalamocortical loop that elucidate how the transitions of beta synchrony from low to high levels might occur in Parkinson’s disease. We provide a new perspective for the effective coupling of the Parkinsonian thalamocortical network, where a fine regulation of temporally different inputs to specific laminae of the motor cortex may underlie the spontaneous and transient variability in oscillatory neural activity in the beta band across the circuit.

## Supplementary

Throughout this study, we were interested in synaptic modulation in the TC loop during beta enhancement in spectral data from 6-OHDA-lesioned rats. This was motivated by recent PD literature defining beta oscillations as intermittent events where power waxes and wanes within a time window of a second. To study the dynamic properties of this parkinsonian biomarker, many of the studies in the field focus on a binary characterization of the beta signal in the PD clinical context while using the envelope of the beta filtered signal and an arbitrary threshold of its mean to divide beta into physiological epochs and hypothetically pathological epochs (Feingold et al., 2015; Sherman et al., 2016; Tinkhauser et al., 2017; Little et al., 2013). Some studies have gone further and suggest that the duration of such high beta events is positively correlated with PD motor impairments.

In CSD-DCM, conditions are effectively spectral densities derived from segments of the data one is interested in studying and as such three considerations must be taken into account: 1) the oscillatory activity undergoing a PSD analysis must have a duration of at least two cycles of its frequency (approximately more than 130 msec for beta frequencies); 2) conditions must consistently show different power levels across subjects and 3) PSD analysis does not account for the time evolution of beta power (does not distinguish segments with short vs. long bursts).

Extracting 500 msec segments (>130msec) with extremely high/low sustained levels of beta power (area below the envelope) allowed us to obtain non-overlapping conditions across subjects with a satisfactory frequency resolution and different functional features: low beta hypothetically accounting for a physiological coupling state and high beta possibly accounting for a state with a higher probability of pathological coupling state.

### Ad hoc beta burst analysis within conditions

Although we were unable to model beta bursts using CSD-DCM, we analyzed the burst density in both low and high beta conditions a posteriori. Making use of the 75% percentile of the mean envelope as a threshold, the mean number of bursts in the LB condition (across trials and subjects) was 0.9 ± 0.3 and those bursts had the mean duration of 195 ± 106 msec.

The mean number of bursts in the HB condition was 3.9 ± 0.7 and had the mean duration of 772 ± 188 msec. As expected, not only HB conditions show a higher density of beta bursts than LB conditions (approximately 4:1) but also its bursts show a longer duration. In both conditions, some trials showed incomplete bursts that were still included in the computation of average number of bursts and averaged duration of bursts per condition.

**Fig.S.1.**
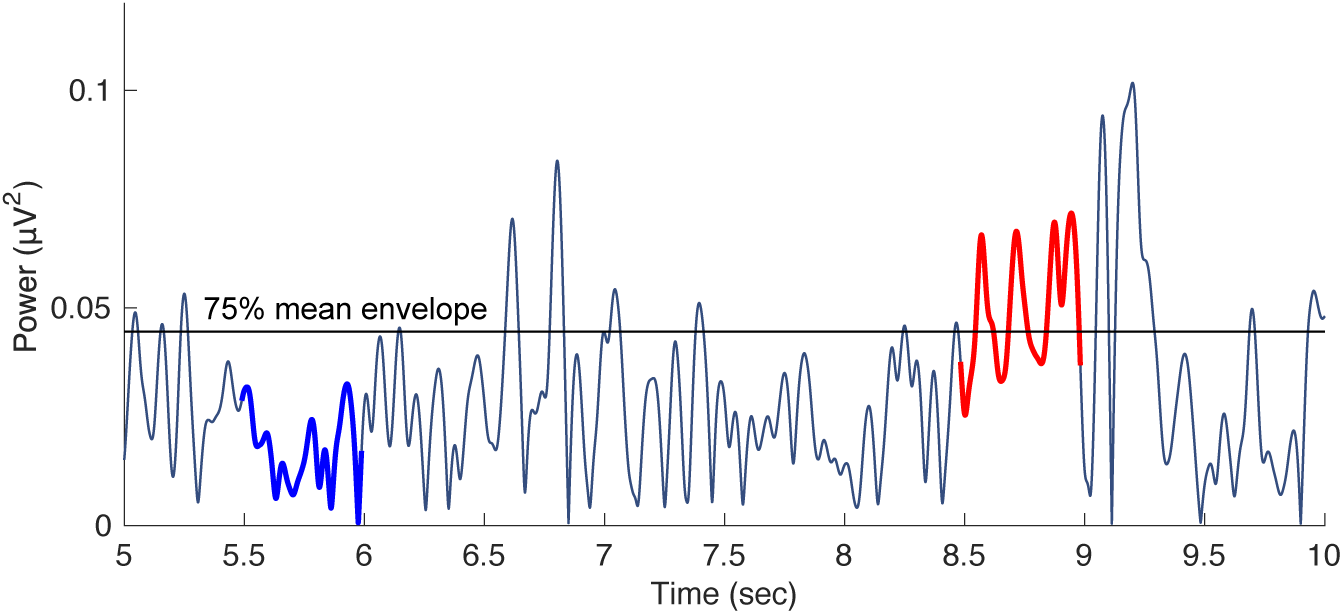
Beta conditions and beta bursts. Example of Beta bursts analysis in the Low and High beta conditions. In this example we can observe how low beta (in blue) and high beta (in red) have different levels of power and present different number of beta bursts. LB with no beta burst and HB with 3 beta bursts.

**Fig.S.2.**
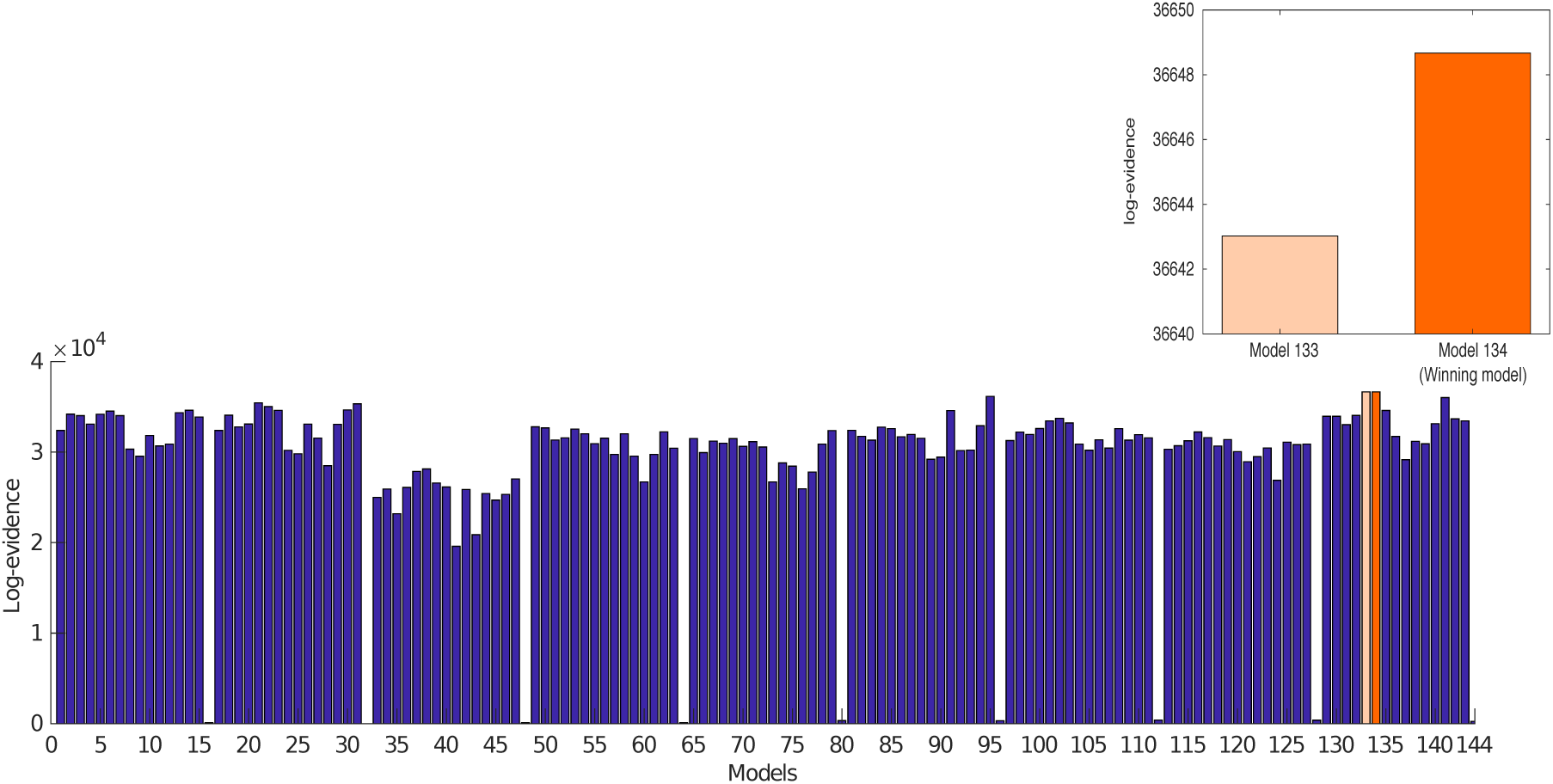
FFX-BMC analysis of the 144 models and winning model. From the 144 models (9 architectures x 16 modulatory configurations), model number 134 (orange) was the one with the higher log-evidence value (3.6649×10^4^) showing a difference of approximately 6 from model 133 (beige) whose log-evidence was the second highest (3.6643×10^4^). Second most plausible model being the model with changes in effective connectivity among the following subpopulations: superficial and deep pyramidal cells reciprocally, relay and reticular cells reciprocally, relay cells to deep pyramidal cells and inhibitory interneurons and deep.

From the averaged posterior estimations on parameters derived from PEB (Fig.6), we computed the net activity of each subpopulation (if net-excited or net-inhibited) during both LB and HB conditions. The same procedure has been applied to the remaining models and can be found in the Supplementary section in (Fig.S.4). This analysis was performed by weighting the combined coupling strength from different efferent subpopulations to an afferent subpopulation; i.e., by using the fixed parameter values of intrinsic and extrinsic connections and the posterior estimations of A/G parameters in the baseline condition and A/G plus B parameters in the HB condition. As observed in the bar plot below (Fig.S.3), the net-activity of our neural mass model presents similar results in both conditions: SP, DP, II and REL receiving a stronger excitatory input than inhibitory (net-excited) and MP and RET receiving a stronger inhibitory input than excitatory (net-inhibited). It is worth noting that to induce a transient beta increase all subpopulations increased their baseline level of activation, i.e., subpopulations that were net-excited in the low condition became more excited in the high condition and subpopulations that were net-inhibited in the low condition suffered further inhibition the high condition. This effect is most predominantly observed in SP, followed by MP and DP.

**Figure.S.3.**
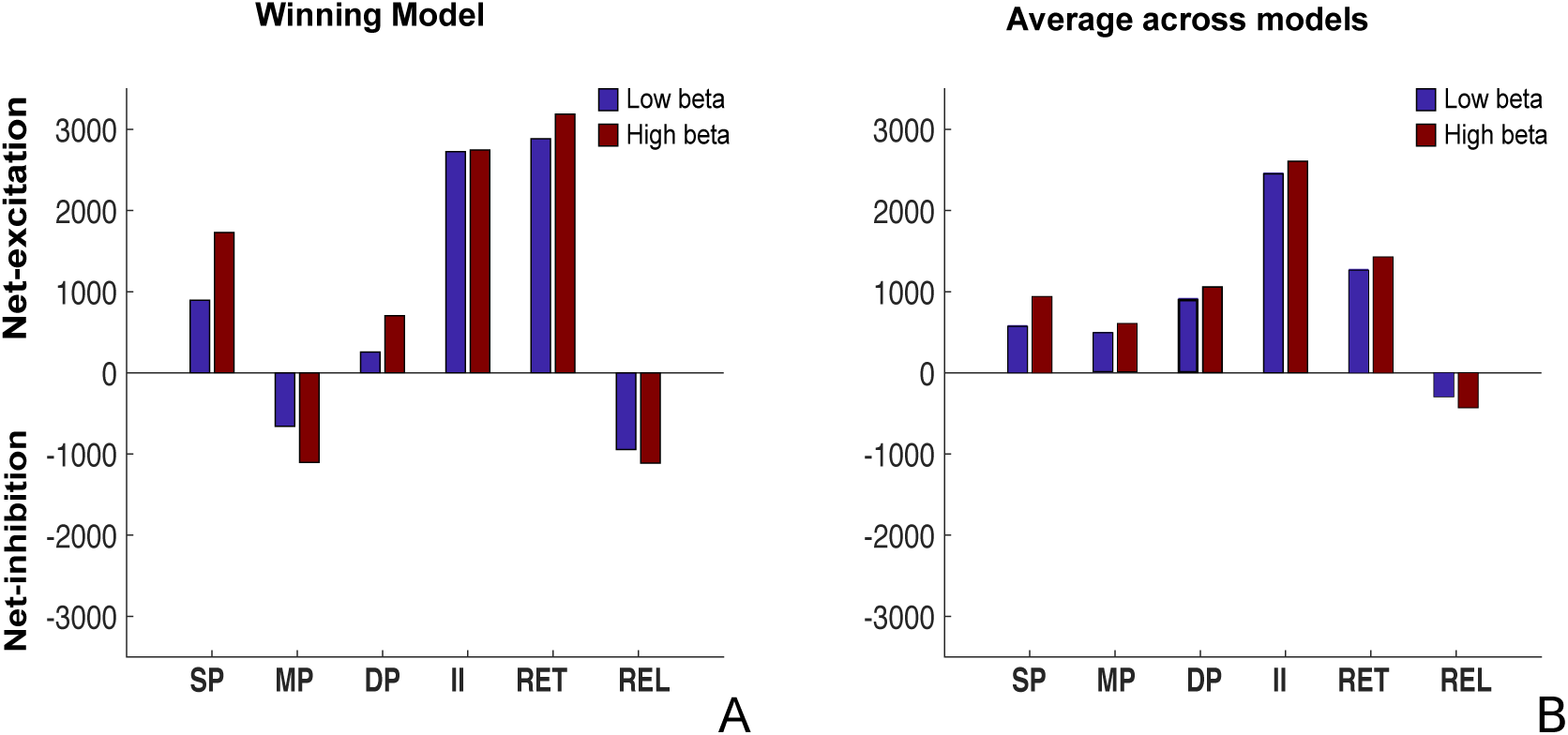
Net-activity state of subpopulations. Overall activation state of neural mass model subpopulations in the different conditions. Bar plot of net-activity of subpopulations in the LB condition (blue) and high beta condition (red). Of interest is the comparison of net-activation levels of each subpopulation in the different conditions and consequently noting that SP is the subpopulation that shows a higher alteration in net-activity (followed by MP and DP).

**Fig.S.4.**
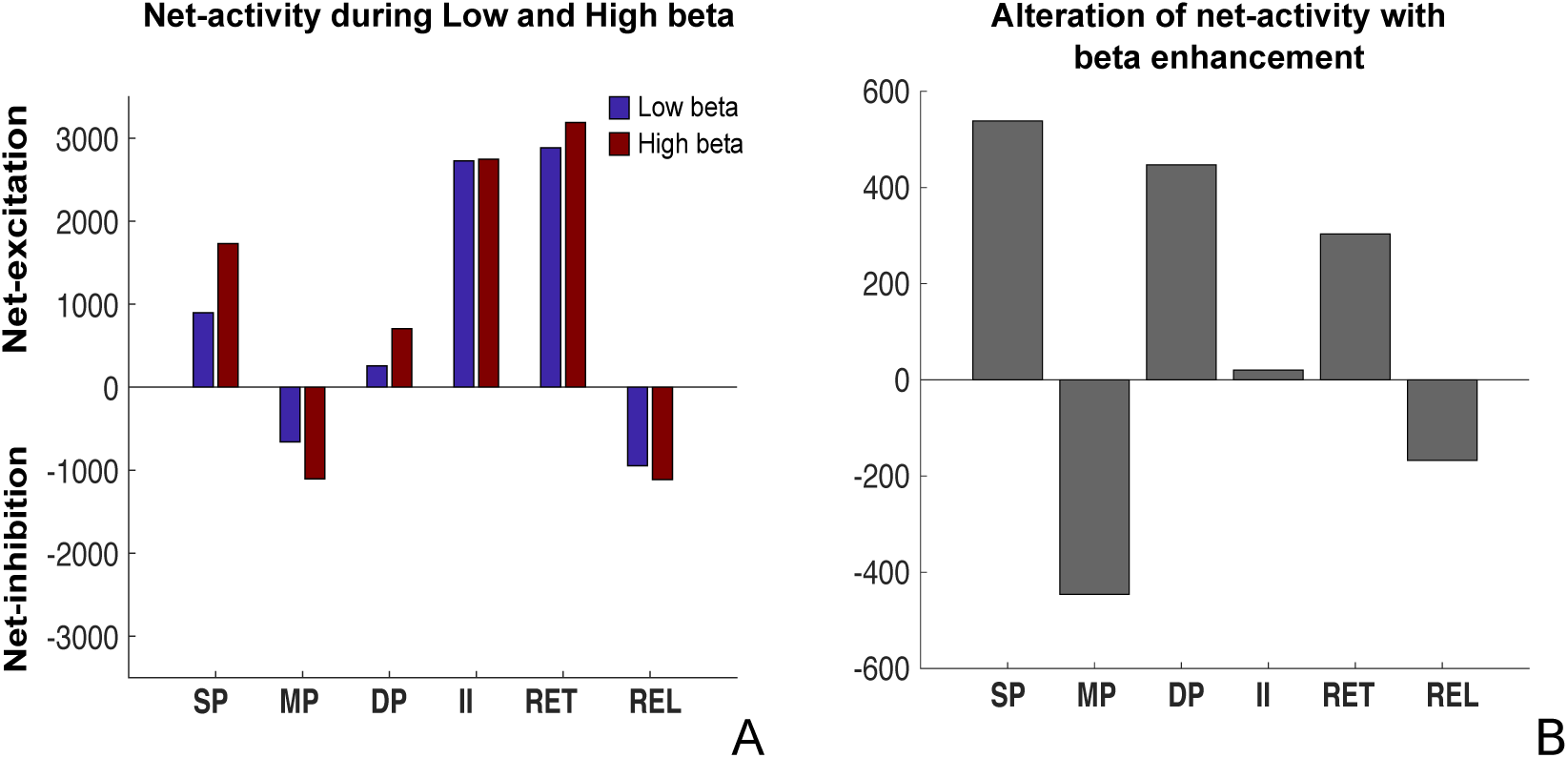
Net-activty state of subpopulations at low and high beta.Comparing the net-excitability pattern across the winning generative model and the average of all models (n=144), we can observe a similar directionality of effects across populations except for the middle pyramidal subpopulation. Also, the mechanism of the created state space (independently of the modulatory pathways) to achieve the spectral transition in our data is consistent between the two plot-an intensification of the baseline level of activation.

## Acknowledgements

This work was supported by studentships (BRT00040) and research funding (MR/R020418/1) from the Medical Research Council (MRC). AS was supported by the MRC (award MC_UU_12024/1). PJM was supported by the MRC (awards UU138197109, MC_UU_12020/5 and MC_UU_12024/2) and Parkinson’s UK (grant G-0806). BW received funding from the European Union’s Horizon 2020 research and innovation programme under the Marie Sklodowska-Curie grant agreement No 795866. TP is supported by the Rosetrees Trust (173346). KJF is a Wellcome Principal Research Fellow (Ref: 088130/Z/09/Z). We thank Dr. N. Mallet for acquiring some of the primary data sets in rodents.

